# Spatial Organization of Gas Vesicles is Governed by Phase-separable GvpU

**DOI:** 10.1101/2023.06.01.543273

**Authors:** Zongru Li, Qionghua Shen, Yifan Dai, Andrew P. Anderson, Manuel Iburg, Richard Lin, Brandon Zimmer, Matthew D. Meyer, Lingchong You, Ashutosh Chilkoti, George J. Lu

## Abstract

Gas vesicles (GVs) are microbial protein organelles that support cellular buoyancy, and the recent engineering of GVs has led to multiple applications including reporter gene imaging, acoustic control, and payload delivery. GVs often cluster into a honeycomb pattern to minimize their occupancy of cytosolic space; however, the molecular mechanism behind this process and its influence on cellular physiology remain unknown. Here, we identified GvpU as the protein governing this process. GvpU-mediated clustering is selective to the genotype of GVs, allowing the design of GV variants with genetically encodable clustering states. Furthermore, we uncovered that the clustering is modulated by phase transition behaviors encoded in the intrinsically disordered region of GvpU through a balanced contribution of acidic and aromatic residues, and such phase transition can directly modulate cellular fitness. Collectively, our findings elucidate the protein player, molecular mechanism, and functional roles of GV clustering, and its programmability for biomedical applications.

---

Organelles are critical components of cells. They often adopt unique spatial organization inside cells, which may facilitate more efficient metabolic flux or render better biophysical properties than an unorganized ensemble^1^. For example, magnetosomes are aligned in a chain configuration inside magnetotactic bacteria to maximize the magnetic moment^3^, and the even distribution of carboxysomes along the axis of a bacterium ensures their segregation during cell division and the survival of daughter cells^4^. Intriguing molecular mechanisms and protein players have been discovered that encode the spatial organizations of these microbial organelles^5-7^. Similarly, gas vesicles (GVs) are a unique class of gas-filled hollow protein organelles inside bacteria and archaea, which use them as flotation devices to compete for the surface of the water to maximize photosynthesis^8^. For cells to achieve a density lighter than water, close to 10% of the cytosolic space needs to be filled by GVs, and thus it has been hypothesized that evolutionary pressure is placed to optimize the efficiency of spatially organizing these protein nanostructures, so that cytosolic space can be freed up for other cellular processes. Indeed, GVs were often observed as uniaxially aligned, honeycomb-patterned bundles in the native cyanobacterial hosts (**Fig. 1a**), and for cylindrical objects, this packing represents the theoretically optimal strategy to minimize the occupancy of cytosolic space with a local air content close to 90%^9^. Despite the recognition of such unique spatial organization of GVs for decades, little is known to date about the protein component or the molecular mechanism behind this process. Besides fundamental biology, the interest in GVs has grown substantially due to the recent development of a wide range of biomedical applications, such as reporter gene and biosensor imaging by ultrasound, MRI, and optical methods^10-18^, cellular control in deep tissues^19, 20^, and gas delivery for photodynamic and sonodynamic therapies^21-23^. To this end, the spatial organization (the clustering) of GVs plays an important role in determining the imaging contrast, the biodistribution, and the cellular interactions of these nanostructures. For example, clustered GVs show a large (more than 10-fold) increase in MRI contrast compared to monomeric GVs^13^. Thus, it is critical to identify the protein components that govern the clustering state of GVs, which will open up avenues to design genetically encodable methods to modulate their imaging contrast in living cells and design clustering-based biosensors.

**Fig. 1.**
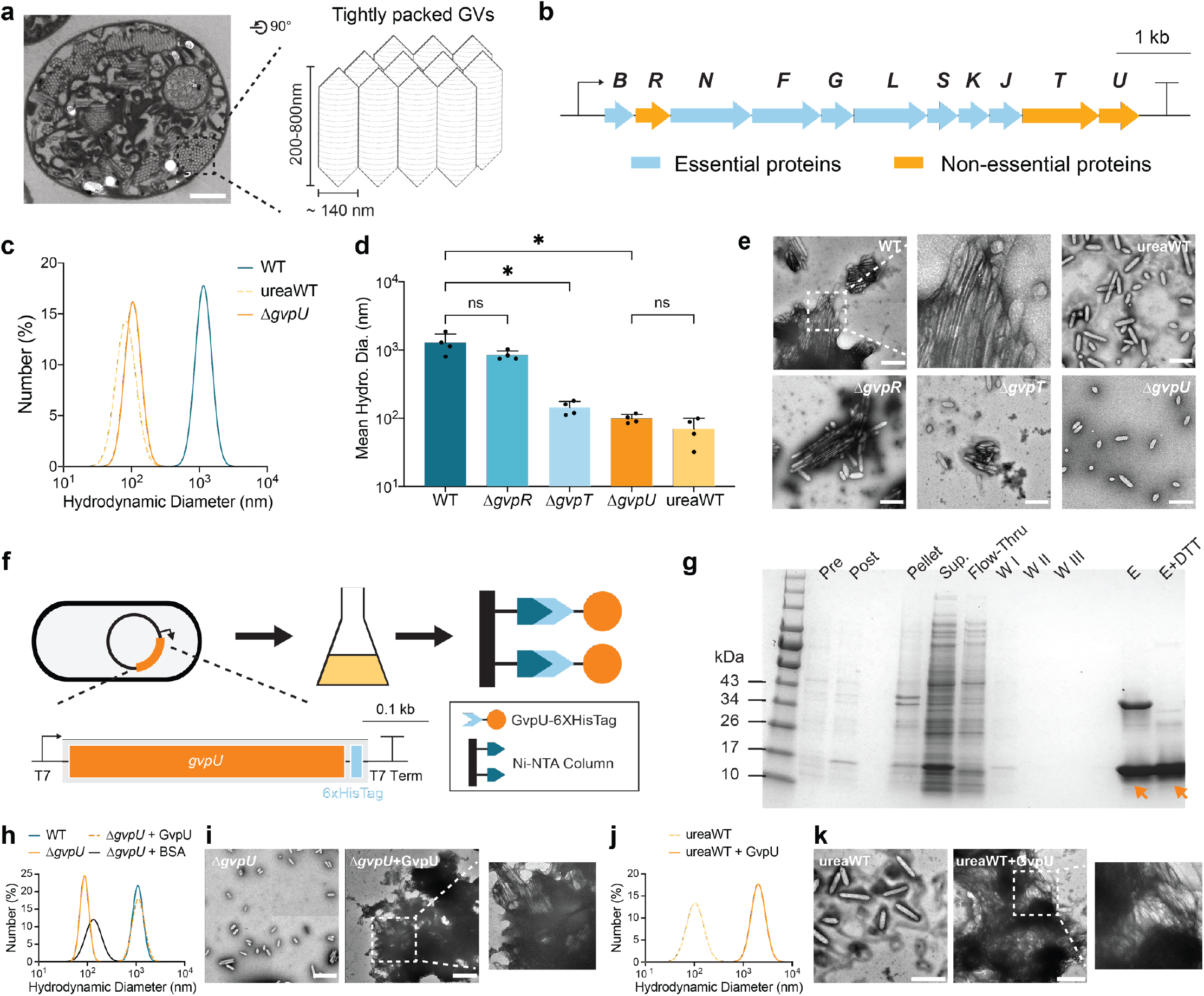
Gene knockout screening determines GvpU is essential for the clustering of GVs and purified GvpU reconstitutes the GV cluster. **a**, Thin-sectioned TEM image of an *Anabaena flos-aquae* cell containing hexagonally packed, clustered GVs, and the schematics drawing of clustered GV protein nanostructures. **b**, The architecture of the GV operon from *Bacillus megaterium* (Mega GVs). Non-essential genes were labeled orange. **c, d, e**, Representative DLS measurement of the hydrodynamic diameter, mean hydrodynamic diameter (n = 4 biological replicates), and representative TEM images of GV variants. WT, wildtype Mega GVs; ureaWT, wildtype Mega GVs after unclustering by 6M urea treatment; *ΔgvpR, ΔgvpT, ΔgvpU*, wildtype Mega GVs without GvpR, GvpT or GvpU, respectively. Error bars represent mean ± standard deviation (STDEV). ns, not significant; *p value < 0.05. Scale bar = 500 nm. **f, g**, Schematic and SDS-PAGE results of the recombinant protein expression and affinity chromatography purification of GvpU::6xHisTag. The protein bands corresponding to GvpU::6xHisTag were labeled with arrows. Pre: pre-induction; Post: post-induction; Pellet and Sup.: pellet and supernatant after cell lysis and centrifugation; Flow-Thru: flow-through of Ni-NTA chromatography; W I, W II, and W III: the washing fractions of Ni-NTA chromatography; E: elution fractions of Ni-NTA-chromatography; E + DTT: elution fractions mixed with DTT to reduce dimer proteins to monomers. **h, i**, DLS measurement and TEM images of wildtype Mega GVs (WT), *ΔgvpU* GVs (*ΔgvpU*), *ΔgvpU* GVs mixed with purified GvpU proteins (*ΔgvpU* + GvpU), and *ΔgvpU* GVs mixed with pure bovine serum albumin proteins as a sham control (*ΔgvpU* + BSA) (n = 3, biological replicates; scale bar = 500 nm). **j, k**, DLS measurement and TEM images of urea-treated GVs (ureaWT) and urea-treated GVs mixed with purified GvpU proteins (ureaWT+GvpU) (n = 3, biological replicates; scale bar = 500 nm).

In this study, to uncover the molecular components and the underlying mechanisms of GV clustering, we started with the natively clustered GVs from *Bacillus megaterium* (Mega GVs) and hypothesized that one of the non-essential genes in the GV operon may encode the clustering state. Gene knockout screening identified GvpU as the protein responsible for the clustering, which was confirmed by *in vitro* reconstitution of the clustering state by purified GvpU. We then discovered through modeling and experiments that GvpU selectively binds to the C-terminal motif of the GV major shell protein. Thus, GvpU-mediated clustering is specific to the genotype of GVs and is orthogonal to the binding of the minor shell protein, GvpC. Based on these findings, we developed a method to genetically encode the clustering state through engineering GV major shell protein. To further determine the biophysical mechanisms of GvpU-mediated clustering, we discovered that GvpU is a protein that can undergo phase separation, and the phase transition is dictated by the intrinsically disordered regions (IDR) of the GvpU through a balance between acidic and hydrophobic residues. A sequence-ensemble-function ternary relationship is established based on the compositions of the IDR, providing evidence on how phase transition can directly benefit cellular physiology. Our findings provide a new mechanism to understand how bacteria organize cellular space to regulate cellular fitness and open up a new dimension of GV engineering by controlling the high-order clustering.

## Gene knockout screening determines that GvpU is essential for the clustering of GVs

To hunt for the protein that governs the clustering of GVs, we selected the GV operon from *Bacillus megaterium* (Mega GVs) for three reasons. First, this operon supports a high level of heterologous expression and assembly of GVs in *Escherichia coli*^24^; secondly, the essential genes for the formation of Mega GVs have been determined before^11^; and thirdly, Mega GVs remain clustered after purification from *E. coli*. Thus, we hypothesized that the protein controlling the clustering state of GVs would: (i) reside inside this GV operon, because bacterial genes responsible for the same function are typically organized into the same operon, and (ii) be one of the non-essential genes, because the clustering of GVs may occur independently from the assembly of the GV shell. To this end, the Mega GV operon consists of 11 genes (**Fig. 1b**), and *gvpBNFGLSKJU* were identified as essential genes^11^, which leaves *gvpR* and *gvpT* as non-essential genes. We compared this list to an earlier study on the GV operon from *Halobacterium salinarum* (Halo GVs), which identified a slightly different set of 8 essential genes, *gvpAOFGJKLM*^25^. As *gvpA* and *gvpM* in *H. salinarum* are homologous to *gvpB* and *gvpS* in *B. megaterium*, respectively, we inferred from this study that *gvpN* and *gvpU* could be additional non-essential genes. Combining these two pieces of information, we set out to conduct individual gene knockout screening on 4 candidates, *gvpR, gvpN, gvpT*, and *gvpU*.

The experimental workflow leveraged the recently established pipeline of GV preparation and characterization, including *E. coli* expression, centrifugally assisted flotation, hydrostatic collapse measurement, dynamic light scattering (DLS), and transmission electron microscopy (TEM)^24^. We observed the successful formation of GVs in 3 out of the 4 single-deletion mutants except for *ΔgvpN*, which produced substantially fewer GVs and supported the conclusion that *gvpN* is an essential gene. Following the purification, the clustering state of GV was determined in solution by DLS and TEM using the wildtype Mega GVs as the control for the clustered form (**Fig. 1c, d, e**). To prepare a control sample for the unclustered form, we followed the established protocol of treating the wildtype Mega GVs with 6M urea to strip off surface proteins and uncluster while preserving the integrity of the protein shell^24^. The resulting clustered GVs typically have a hydrodynamic diameter of ∼1000 nm in solution, while the unclustered GVs have a hydrodynamic diameter of ∼100 nm, providing a large dynamic range for DLS to distinguish the clustering states of the GVs being tested. Our results showed that *ΔgvpR* remained clustered. While both *ΔgvpT and ΔgvpU* GVs showed a reduction in the hydrodynamic diameter as compared to wildtype Mega GVs, bundles of clustered GVs remained visible in TEM images of *ΔgvpT* GVs, and only *ΔgvpU* GVs were fully unclustered. Thus, the screening identified *gvpU* as the top candidate gene for governing the clustering of GVs.

## Purified GvpU reconstitutes the GV cluster

To confirm that *gvpU* is the protein governing the clustering of GVs, we next asked the question of whether the presence of GvpU is sufficient to reconstitute the clustering of the monomeric GVs. To address this question, we needed to first develop an expression and purification protocol for GvpU. A poly-histidine tag was appended to the C-terminus of *gvpU*, and the fusion proteins were induced for expression in *E. coli*, followed by purification by immobilized metal affinity chromatography (**Fig. 1f)**. Sufficiently pure proteins corresponding to the molecular weight of GvpU were obtained by this method (**Fig. 1g)**, and mass spectrometry confirmed its identity as GvpU (Supplementary Fig. 1). Next, we added the purified GvpU protein to *ΔgvpU* GVs, and both DLS and TEM demonstrated the formation of clustered GVs after brief incubation (**Fig. 1h, i**). These results confirm that GvpU is sufficient to confer the clustering function of GVs. To further rule out the participation of other surface-bound proteins in the clustering of GVs, we repeated the *in vitro* reconstitution using 6M urea-treated Mega GVs, and the subsequent DLS and TEM analysis showed similar results, confirming that GvpU alone is sufficient to induce the clustering (**Fig. 1j, k**).

## GvpU-mediated clustering is selective to the genotype of the GV major shell protein

Next, we set out to test the hypothesis that GvpU-mediated clustering is specific to the genotype of GVs. We chose to evaluate the ability of GvpU to mediate the clustering of another genotype of major shell proteins, GvpA, from *Anabaena flos-aquae*. GvpA can be successfully assembled into GVs in *E. coli* by the same set of assembly factor proteins with the only difference being two copies of *gvpA* are needed to boost their expression (**Fig. 2a**)^10^. Experimentally, we constructed the GvpA-based GV operon with *gvpU* (*A*_WT_) and without *gvpU* (*AΔgvpU*), and the resulting DLS and TEM showed that both genotypes displayed a similar hydrodynamic diameter of ∼100 nm (**Fig. 2b, c, d**). In contrast to GvpB-based *ΔgvpU* GVs (*BΔgvpU*, **Fig. 1h, i**), adding purified GvpU back to the *AΔgvpU* variant did not result in the clustering of these GVs. Furthermore, we evaluated if GvpU directly binds GV nanostructures. To this end, purified GvpU was incubated with *AΔgvpU* or *BΔgvpU* GVs for binding, followed by centrifugally assisted flotation (**Fig. 2e**). As expected, SDS-PAGE revealed that GvpU was only present when incubated with *BΔgvpU* (**Fig. 2f**). Thus, these results establish that GvpU specifically interacts with the major shell protein, GvpB, but not with another major shell protein such as GvpA.

**Fig. 2.**
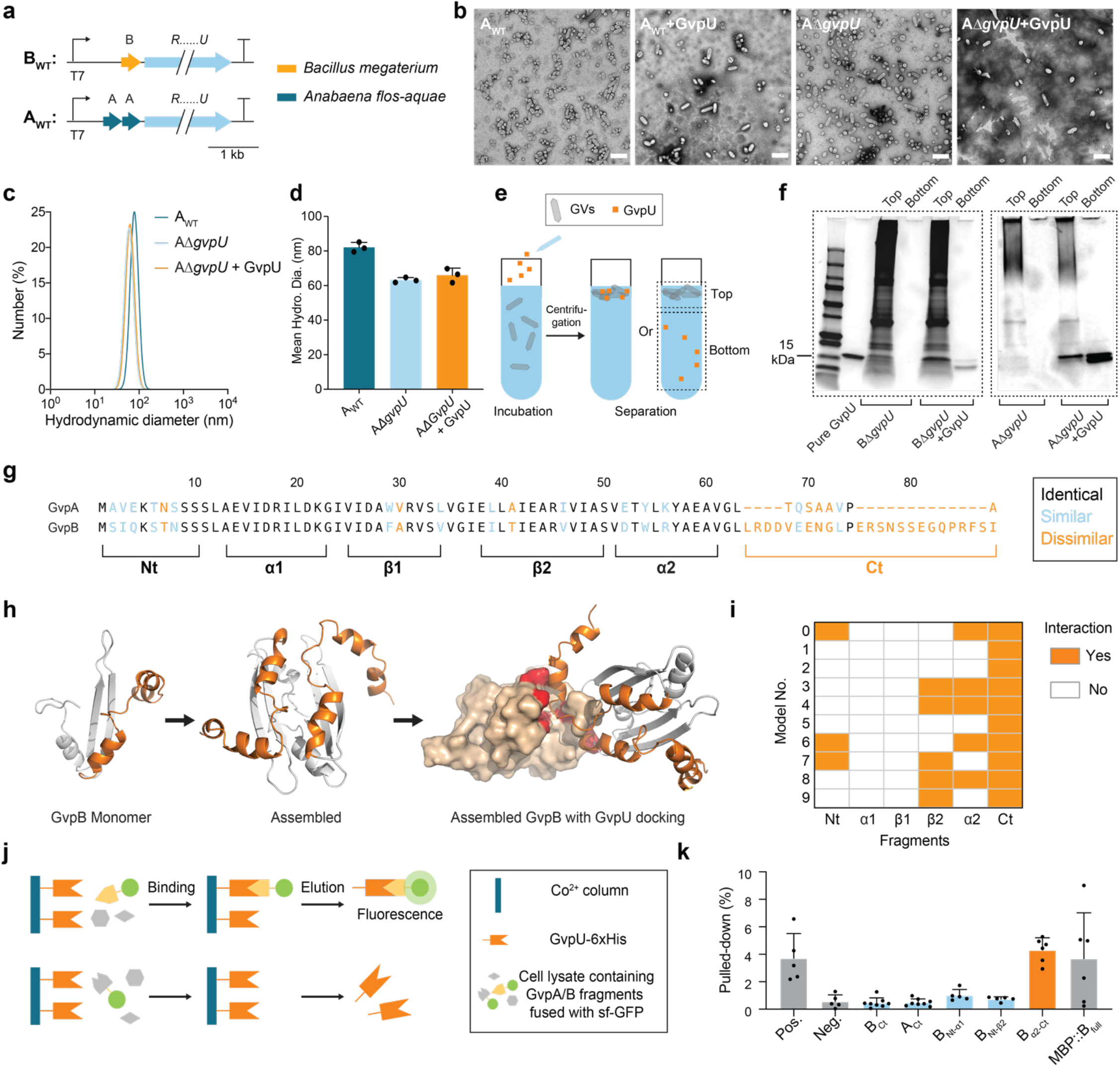
GvpU-mediated clustering is selective to the genotype of the GV major shell protein and the C-terminal region of the GV major shell protein is the binding site for GvpU. **a**, The architecture of the gene clusters that support the assembly of two genotypes of major shell proteins in *E. coli*. A and B are the major shell proteins, *gvpA* and *gvpB*, respectively, and R…U is the assembly factor protein cassette consisting of *gvpRNFGLSKJU* that supports the assembly of the major shell proteins into GV nanostructures. **b, c**, Representative TEM images and DLS measurements of GvpA-based GVs with GvpU (A_WT_) and without GvpU (*AΔgvpU*) and GvpB-based GVs with (B_WT_) and without GvpU (*BΔgvpU*). Scale bar = 500nm. n = 3 biological replicates for DLS measurements. **e, f**, Schematic and SDS-PAGE results of the experiment to assess the direct binding of GvpU to the floating GV nanostructures. After the addition of purified GvpU, incubation, and centrifugation, samples from the top layer that contains both GVs and the media (Top) and the bottom layer that contains only the media (Bottom) were run on SDS-PAGE. Arrows indicate the location of the GvpU proteins on SDS-PAGE. **g**, Sequence alignment of GvpA and GvpB. The secondary structure can be divided into the N- and C-terminal regions (Nt and Ct), two α-helices α1 and α2, and two β-strands β1 and β2. **h**, Representative simulated docking of GvpB and GvpU. Two Rosetta-modelled GvpB monomers were assembled into a dimer, and the top 10 models of GvpU docking to the GvpB dimer were selected for analysis. The α2-Ct segment of GvpB was labeled in orange, and the docking sites on GvpU were labeled in red. **i**, Heat map of GvpB-GvpU interactions in the top 10 docking models. The segments of GvpB that have contact with GvpU are labeled in orange. **j, k**, Schematic of *in vitro* pull-down assay and normalized pulled-down percentage of each bait-prey pair. GvpU was first immobilized on Co^2+^ affinity column, followed by the flowthrough of the cell lysates containing the fusion protein of GvpB and GvpA with sfGFP. The fluorescence intensity of sfGFP was quantified after the elution of the fusion protein (n ≥ 5 biological replicates). The positive control (Pos.) uses the covalent interaction between SpyCatcher and sfGFP::SpyTag^2^, and the negative control (Neg.) uses sfGFP without the fusion of the fragments from GvpA or GvpB. Another positive control, MBP::B_full_, measures the interaction of full-length GvpB with GvpU. Due to the insolubility of full-length GvpB, the maltose-binding protein was fused to GvpB (MBP::B_full_) to enhance the solubility, and this construct was further fused to sfGFP. Error bars represent mean ± STDEV. Error bars represent mean ± STDEV for n = 3 biological replicates. **** p < 0.0001.

## The C-terminal region of the GV major shell protein is the binding site for GvpU

Next, we seek to determine the structural motifs of the GV major shell protein that mediate their interaction with GvpU. To this end, the secondary structure of the major shell proteins consists of an α-ß-ß-α topology flanked by the less conserved N-and C-terminal tails (Nt and Ct) (**Fig. 2g**). We first performed computational docking of GvpU with GvpB. A dimer of GvpB was chosen to better mimic the assembled GV nanostructures, and the analysis of ClusPro’s top 10 docking models revealed that the C-terminal motifs (ß2, α2, and Ct) have the highest chance of interaction with GvpU (**Fig. 2h, i**). Guided by this prediction, we performed a set of *in vitro* pull-down assays on cell lysates containing various fragments of GvpB and GvpA using an affinity column with immobilized GvpU (**Fig. 2j**). Based on the observed similarities and dissimilarities between GvpA and GvpB, a series of fragments were generated (**Fig. 2g & Supplementary Table 1**). All fragments were fused with superfolder GFP (sfGFP) to facilitate their expression and quantification. Interestingly, the C-terminal tail of GvpB alone (B_ct_) does not confer strong binding to GvpU; instead, including the second α-helix (B_α2-Ct_) rescued the binding of the fragment to the same strength as that of the full-length GvpB. Complementary to this finding, various constructs of the N-terminal region of GvpB showed minimal interactions (**Fig. 2k**). Together, these results established that GvpU specifically interacts with the second α-helix and the C-terminal tail of GvpB, which is in good agreement with the docking simulations.

## The minor shell protein GvpC promiscuously binds to Mega GVs without interfering with clustering

A minor shell protein of GVs, GvpC, is present in various genotypes of GVs such as those of *A. flos-aqua* (Ana GVs) and *Halobacterium salinarum* (Halo GVs), and GvpC usually binds to the outer surface of the major shell proteins (**Fig. 3a**)^26, 27^. Here, we question the modularity of GvpC and GvpU binding and whether their binding will interfere with each other. To approach this question, we first constructed several GV variants, A2C, BC, and B2C, in which the names denote the copy number of the major and minor shell protein genes (**Fig. 3b**). To our knowledge, it has not been shown before whether GvpB from *B. megaterium* has enough conserved motif to allow the binding of GvpC from *A. flos-aqua*, and thus it is interesting to see that GvpC were present in all of these GV variants we tested (**Fig. 3c**), which established for the first time that the GvpB-based GVs can promiscuously accommodate the surface binding of GvpC from *A. flos-aqua*. This conclusion was further corroborated by the observed increase of the hydrostatic collapse pressure of BC and B2C from that of wildtype GvpB-based GVs, consistent with the notion that GvpC strengthens the mechanical properties of GVs (**Fig. 3d, e**)^9^. Lastly, we set out to evaluate whether the binding of GvpC would interfere with the GvpU-mediated clustering of these GVs. We carried out TEM and DLS measurements of these GVs variants, which revealed that the clustering states remained the same regardless of the presence of GvpC (**Fig. 3f, g**). We also isolated Δ*gvpU* variants of B2C GVs, which remained unclustered, indicating that GvpC does not induce clustering (**Supplementary Fig. 2**).

**Fig. 3.**
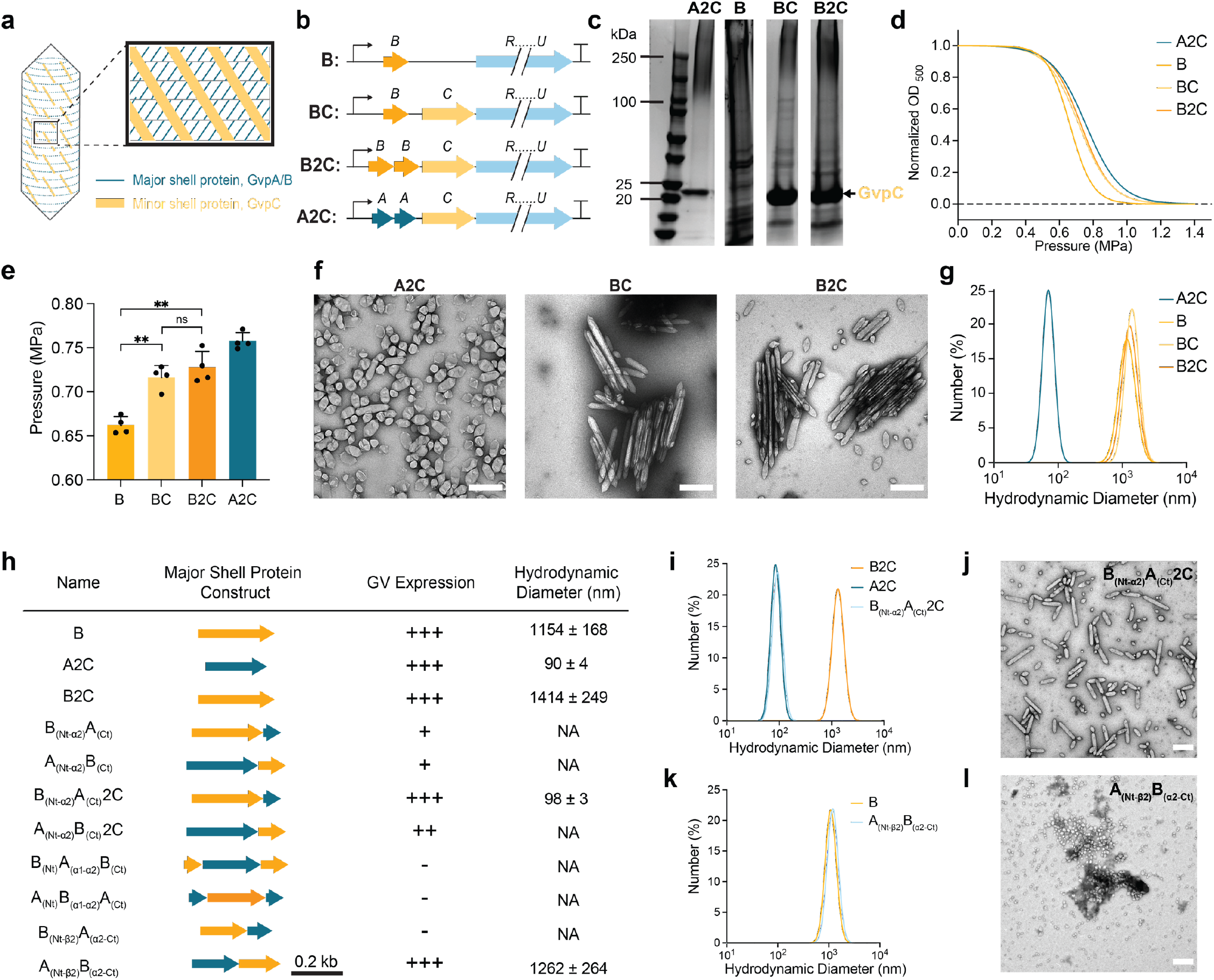
The minor shell protein GvpC promiscuously binds to Mega GVs without interfering with clustering, and the engineering of the major shell protein to toggle between clustered and unclustered states. **a**, Schematic diagram of GV surface, on which the minor shell protein GvpC wraps on the outside of tightly packed lattice of the major shell protein, GvpA/B. **b**, The genetic architectures of B, BC, B2C, and A2C GV variants, of which the names denote the genotype and the copy number of the major and minor shell protein genes, respectively. **c**, SDS-PAGE result of GV variants after purification. The band corresponding to GvpC is labeled in yellow. **d**, Representative hydrostatic collapse curves and normalized optical density at 500nm (OD_500_) quantifies the fraction of the intact, gas-containing GVs before being collapsed by a step-wise increase of hydrostatic pressure. Curves represent fits of the data using the Boltzmann sigmoid function, and the characteristic V50 values are used as the midpoint values. **e**, The mean midpoint collapse pressure as a measurement of the mechanical properties of GV variants. Error bars represent mean ± STDEV for n = 4 biological replicates. ns, not significant; **p value < 0.01. **f, g**, Representative TEM images and DLS measurements of the GV variants. n = 4 biological replicates; scale bar = 500 nm. **h**, A table of the designed major shell protein constructs, the characterization of GV formation in *E. coli*, and the determination of the clustering state *via* the measurement of hydrodynamic diameter. Orange and cyan represent *gvpB* and *gvpA* gene segments, respectively. For GV expression, single positive signs (+) represent GVs that can be observed by naked eyes after cell lysis; double signs (++) indicate that the amount of GVs was enough for TEM sample preparation, and triple signs (+++) indicate the number of GVs was enough for both TEM and DLS characterizations; and the negative sign (-) represents that GVs were not detectable. For hydrodynamic diameter, n = 3 biological replicates were measured for each GV variant, and the mean and STDEV were reported. **i, k**, Representative DLS measurement of B_(Nt-α2)_A_(Ct)_2C compared to the wildtype B2C and A2C constructs, and A_(Nt-β2)_B_(α2-Ct)_ compared to the wildtype B constructs. n = 3 biological replicates. **j, l**, Representative TEM images of B_(Nt-α2)_A_(Ct)_2C and A_(Nt-β2)_B_(α2-Ct)_ GVs. Scale bar = 500 nm.

## Engineering the major shell protein to toggle between clustered and unclustered states

Having a modular motif on GV shell proteins that can toggle GVs between the clustered and unclustered states would provide a powerful method to engineer GVs for biological applications. For example, multiplexed imaging of several GV variants would enable the simultaneous tracking of multiple biological processes, analogous to that achieved by multi-color fluorescent proteins but in deeper tissues, and achieving this goal would require the design of GV variants that can orthogonally adjust parameters important for imaging, such as mechanical properties and clustering states^13^. To this end, GvpA- and GvpB-based GVs possess distinct mechanical properties but predefined clustering states, and thus, it would be highly desirable to develop a method that can modularly toggle the clustering states. In practice, however, mutagenesis of the major shell protein is challenging because of the tight crystalline packing of the shell proteins in the mature GV nanostructures and the involvement of multiple proteins during the assembly process. Since our above results established the region that controls the clustering is close to the C-terminus, which is the most variable region of the protein, we set out to test the possibility of designing new GV shell proteins that retain the ability to assemble into GVs but with toggled clustering states.

We focused on GvpB from *B. megaterium* and GvpA from *A. flos-aquae* and hypothesized that swapping the regions that were identified as the binding site of GvpU would allow us to toggle the clustering state. Additionally, considering the difficulty in assembling mutant GV shell proteins, we planned to screen the inclusion of two copies of the major shell proteins as a method to enhance their expression ^10^ and with and without GvpC in the gene cluster. A list of constructs was screened and, unsurprisingly, the majority of them did not lead to a robust assembly of GVs (**Fig. 3h**). Among the constructs, replacing the C-terminal region of GvpB with that of GvpA (B_Nt-α2_-A_ct_2C) successfully resulted in the loss of the clustering of GVs. On the other hand, Swapping the C-terminal region of GvpA with that of GvpB (A_Nt-α2_-B_ct_ and A_Nt-α2_-B_ct_2C) did not result in an adequate amount of GVs for DLS size quantification, likely due to compromised assembly process. Subsequently, including the second α-helix (A_Nt-β2_-B_α2-Ct_) substantially improved the assembly of GVs, and remarkably, this variant gains the function of clustering as we predicted (**Fig. 3i, j, k, l**). Thus, B_Nt-α2_-A_ct_2C and A_Nt-β2_-B_α2-Ct_ established the possibility of engineering the major shell protein with toggled clustering states.

## GvpU can drive phase separation through homotypic interactions

After establishing the role of GvpU in mediating GV clustering and the engineerability of the shell proteins, we set out to investigate the underlying biophysical mechanism by which GvpU enables GV clustering. The clustering effect or assembly of a high-order network, which is widely observed in the study of patchy colloids^28^, suggests the existence of multivalent interaction sites on the GV, thereby forming a connected network^29^. As the reconstitution of GvpU results in the assembly of a GV network, we hypothesized that the binding of GvpU to GV provides GV with a homotypic self-interaction domain, thereby mediating GV clustering. To this end, we first tested whether GvpU can mediate homotypic interactions. During the purification of GvpU, we indeed observed the reversible formation of turbid mixtures upon a change in the ionic strength (**Fig. 4a**). This phenomenon has been observed in proteins that drive phase transitions and a common feature of such proteins is the existence of intrinsically disordered regions (IDRs)^30-32^. To uncover whether GvpU has such IDRs, we conducted an analysis of GvpU using a meta-predictor^33^, which predicts structural disorder with a consensus artificial neural network. Regions with a significant disorder score were observed between the residues 50-100 and at the C-terminus (**Fig. 4b**). To experimentally test whether GvpU exhibits a phase transition behavior, we examined the turbid mixture of purified GvpU::sfGFP fusion proteins through fluorescence microscopy. We found that the turbid mixtures were the results of spherical condensates and the formation of such mixture is a concentration-dependent process and also salt-sensitive that are consistent with biomolecular phase separation (**Fig. 4c**)^34^. To exclude the effect of sfGFP fusion in driving phase transition, we used a non-fluorescently labeled GvpU and acquired a series of time-lapse phase contrast images, from which we observed that the condensates could coalesce with their neighbors, suggesting a liquid-like behavior (**Fig. 4d**).

**Fig. 4.**
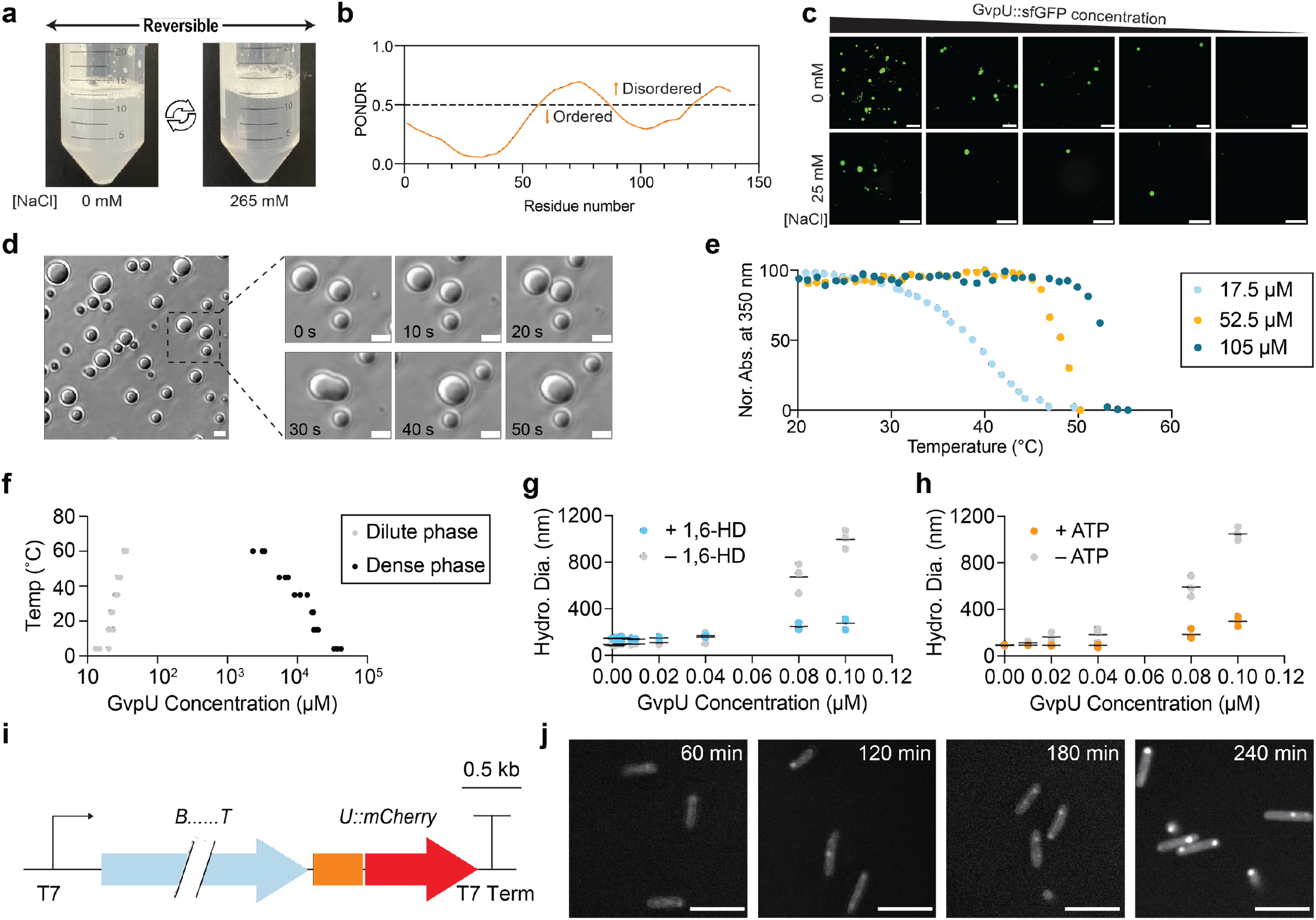
GvpU can drive phase separation through homotypic interactions, and the phase separation of GvpU mediates the clustering of GVs. **a**, Reversible formation of a turbid solution of purified GvpU upon the variation of the ionic strength of the buffer at room temperature. **b**, Prediction of the intrinsically disordered regions of GvpU by meta-predictor, PONDR-FIT. **c**, Fluorescence microscopy images of GvpU::sfGFP in 25 mM Tris-HCl buffer containing either 0 or 25 mM of NaCl at room temperature. The GvpU::sfGFP concentration series are 900, 180, 90, 60, and 45 μM from left to right (scale bar = 5 μm). **d**, Time-lapse phase-contrast images show the dynamic fusion process of GvpU condensates. The solution was prepared with 0.5 mM GvpU in 25 mM Tris-HCl and 250 mM NaCl at room temperature (scale bar = 5 μm). **e**, Temperature-dependent phase transition of GvpU. The turbidity of the solution was monitored at 350 nm as the temperature of the solution was decreased step-wise from 55°C to 20°C. The three different concentrations of the GvpU samples are labeled. **f**, Phase diagram of purified GvpU proteins, which shows a typical UCST phase transition behavior. **g, h**, Hydrodynamic diameter measurement by DLS of GV clusters as a function of GvpU concentration in the absence or presence of chemicals that disrupt phase separation. 1,6-HD, 1,6-hexanediol; ATP, adenosine triphosphate. GvpB-based GVs were held at OD_500_ = 0.2. pH = 7.5 for all the buffer conditions. n = 3 biological replicates. **i**, Genetic architectures GvpB-based GVs, of which the mCherry was fused to the C-terminus of *gvpU*. **j**, Confocal fluorescence microscopy images of *E. coli* cells expressing a modified GvpB-based GV operon, which contains with a *gvpU::mCherry* fusion protein. Red fluorescence images were acquired at different time points post-induction (scale bar = 5 μm).

To further uncover the type of thermodynamic driving forces of GvpU phase transition, we utilized temperature-dependent optical turbidity to analyze the thermos-responsive phase behavior of GvpU under physiological salt conditions^35^. Decreasing the solution temperature resulted in a sharp increase in the turbidity of the solution, and the transition temperature showed a concentration dependency that is characteristic of upper critical solution temperature (UCST) phase behavior^36^ (**Fig. 4e**). The UCST systems, exemplified by polyampholytes such as resilin-like polypeptide^30^, undergo a phase transition commonly through the combinations of electrostatic interactions and pi-based interactions, which is consistent with the fact that the disordered region of GvpU contains a large number of charged residues and aromatic residues. We further constructed the phase diagram of GvpU using a temperature-dependent sedimentation assay (**Fig. 4f**)^32^, and as expected, the binodal curve of GvpU showed a concave down shape that is a signature of UCST phase behavior.

## Phase separation of GvpU mediates the clustering of GVs

Next, to ask whether phase separation of GvpU is the underlying mechanism for GV clustering, we titrated GvpU into unclustered GVs and found a dose-dependence of GvpU on GV clustering and a gradual increase of the size of the GV clusters. This network formation process is similar to the percolation process observed along with phase separation^37^. This phenomenon suggests the existence of a threshold of valences to trigger GV clustering. To further confirm that the phase transition of GvpU is the major driving force for GV clustering, we added 1,6-hexanediol and ATP, which are agents commonly used to perturb phase separation by modulating the weak hydrophobic and electrostatic interactions, respectively^38^. As expected, these reagents significantly inhibited the formation of GV clusters (**Fig. 4g, h**).

We further set out to examine whether we can visualize the GvpU-mediated clustering effect in living cells. To this end, we fused a mCherry fluorescent protein onto the C-terminal of GvpU (**Fig. 4i**) and transformed the GV gene cluster into *E. coli*. After the induction of the GV gene cluster, we tracked the fluorescence signal through confocal fluorescence microscopy at different time points (**Fig. 4j**). A time-dependent, thereby expression level-dependent, puncta formation process was observed, suggesting that the GvpU-mediated clustering effect has a concentration threshold. Furthermore, GvpU::mCherry signal was still notable throughout the cells after the puncta formation, suggesting the existence of a dilute phase of GvpU. This observation confirms that GvpU exists in a phase equilibrium rather than aggregates on GVs, which would be a nucleation process. This observation confirms that GvpU can drive phase transition in living cells.

## Intrinsically disordered region of GvpU drives GV clustering through a balance between acidic and hydrophobic residues

To further understand the correlation between the phase transition of GvpU and GV cluster at the molecular level, we aimed to dissect the dependency of IDR amino acid composition on its clustering function. Based on the established sticker-and-spacer model for understanding IDR sequence-function correlation^36^, we first elected to mutate a sticker-type residue, Phe (F) that contributes to cation-π and π–π interaction, to a spacer type residue, Ser (S) that contributes to the chain solvation (**Fig. 5a**). We observed a significant decrease in the mean hydrodynamic radius of GVs (**Fig. 5b, c**), suggesting that π-based interactions are critical to the assembly of GV clusters. Next, we wondered about the role of Glu (E), which is the only charged residue in the IDR. The mutation of Glu to Ser and Gly transforms the IDR from a polyanion to a non-ionic polymer, which significantly increases the hydrodynamic radius of GV cluster, confirming the importance of hydrophobicity in driving the phase transition of GvpU. This observation further suggests that, first, the heterotypic electrostatic interaction between GvpU and other components on the GV is not the driving force for GV assembly, and secondly, the interspaced acidic residue serves to offset the strong hydrophobicity-driven interactions, thereby achieving an optimum assembly size of the GV cluster. Such an ensemble-function relationship aligns with the observations in human transcription activation domain, in which a balance between hydrophobic and acidic residues is required for strong transcription activation^39^.

**Fig. 5.**
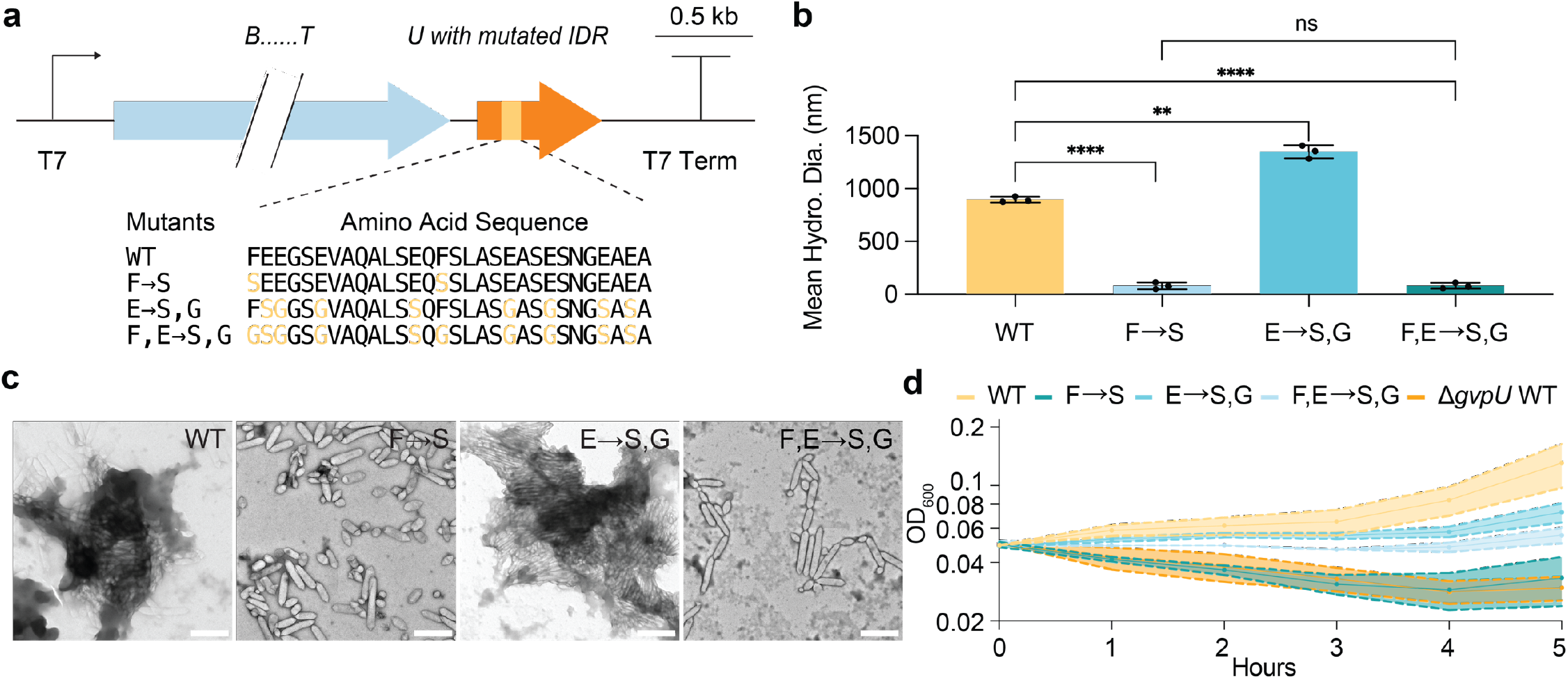
Intrinsically disordered region of GvpU drives GV assembly through a balance between acidic and hydrophobic residues, and IDR composition directly modulates cellular fitness. **a**, Genetic architectures GvpB-based GVs, of which the middle IDR region of *gvpU* was mutated. Mutated residues are labeled in yellow. **b, c**, DLS measurement and TEM images of wildtype GvpB-based GVs (WT), WT GVs with *gvpU* IDR Phe to Ser mutation (F → S), *gvpU* IDR Glu to Ser/Gly mutations (E → S, G), and *gvpU* IDR Phe/Glu to Ser/Gly mutations (F, E → S, G). Scale bar = 500 nm. **d**, Cellular fitness tracking by measuring the optical density of *E. coli* culture at 600 nm (OD_600_). The curves display mean ± STDEV, n = 3 biological replicates. The solid lines represent the mean values, and the shaded area represents one STD above and below the mean. The x-axis shows the time when the data points were collected, and the hours correspond to the time after diluting the GV-expressing cells into fresh media without the inducer.

## IDR composition directly modulates cellular fitness

Lastly, we investigated whether the phase separation-mediated GV cluster correlates with cellular physiology. Since the clustering of GVs has been hypothesized to contribute to freeing up the cytosolic space, we predict that the GvpU-mediated clustering will positively influence cellular fitness. To evaluate this prediction, we implemented differently designed IDR mutants of GvpU to guide the assembly of GVs in *E. coli*. The cellular population growth of *E. coli* was tracked after the formation of GV clusters in a medium without inducer to exclude the cellular burden brought by gene expression. The population containing the wildtype GvpU shows the highest growth recovery, and all three mutants showed significant retardation of growth rate similar to the construct without GvpU (**Fig. 5d**). What is most remarkable is the E → S,G mutant, which enhanced the clustering of GVs *in vitro* (**Fig. 5b**) but such enhancement was shown here to produce a detrimental effect on cellular fitness in the same fashion as the other two mutants. Our results show that an optimal assembly size of the GV cluster is required for cellular fitness, and to our knowledge, this study is one of the first examples of how phase separation can contribute directly to cellular functions^40^, which in our case, is promoting cellular fitness.

## Discussion

To our knowledge, this paper identifies for the first time a protein that governs the clustering of GVs and the related biophysical mechanism of driving the clustering through phase separation, thus filling an important knowledge gap in the spatial organization of this class of protein organelles. Furthermore, we uncovered that cellular fitness is directly correlated with the phase behaviors and the chemical functionality of the protein, which delivers new knowledge on how phase transition and IDR can contribute to cellular physiology.

The intriguing role of biomolecular condensates in driving the clustering of GVs can shed light on a generalizable strategy utilized by cells to organize the subcellular structures^41, 42^. As we have observed, GV clustering is a density-transition process and has a concentration threshold for this transition, which suggests that the GV clustering process is possibly dictated by phase separation^37^. Furthermore, we observed that GvpU could drive phase transition solely, which implicates its role in GV clustering. We uncovered the molecular grammars of the embedded IDR (between the N- and C-terminal structured domains of the GvpU) on driving GV assembly and demonstrated how a balanced electrostatic repulsion and hydrophobic interaction can contribute to the optimum scenario for GV clustering, which directly modulates cellular growth. Based on these observations, we can propose an assembly model of GVs (**Fig. 6a, b, c, d**): GvpU resides on the surface of GVs through domain-specific interactions, and multiple units of GvpU may cover the surface of a single GV, empowering the GV for homotypic interactions through IDR. With sufficient GvpU-covered GVs, IDR of GvpU guides the assembly of the GV clusters. An optimal assembly size is determined by the strength of IDR-sequence-dependent inter-GV interactions, thereby contributing cellular packaging with a possible morphology possessing a minimal interfacial free energy^43^. Such a subcellular spacing strategy directly benefits cellular fitness. In parallel, our study may set the stage for completing the structural understanding of GVs. To this end, two recent cryo-EM studies provided the high-resolution structure of the GV shell proteins for the first time^26, 27^. However, both structures were limited to the first 66 residues, because the C-terminal tails (Asp67-Ile88 of GvpB and Ala67-Ala71 of GvpA) were flexible in the monomeric form of GVs, which were utilized as samples for structure determination. Intriguingly, our study here demonstrated that it is exactly the flexible C-terminal tail that provides the major contribution to the interaction of GvpB with GvpU. Thus, it would be interesting to analyze whether the C-terminal tail of the major shell protein will adopt a defined structure upon the binding of GvpU. If true, studying the major shell protein together with GvpU may be a unique way to determine the full-length structures of the shell protein. Here, we combined an AlphaFold-predicted model of GvpU with the cryo-EM structure of GvpB to illustrate this hypothesis (**Fig. 6e**). Overall, our work may open up avenues for future studies to experimentally determine the structures of the C-terminal tail of the shell protein and GvpU, which will provide a full picture of GV nanostructures with the corresponding inter-particle organization.

**Fig. 6.**
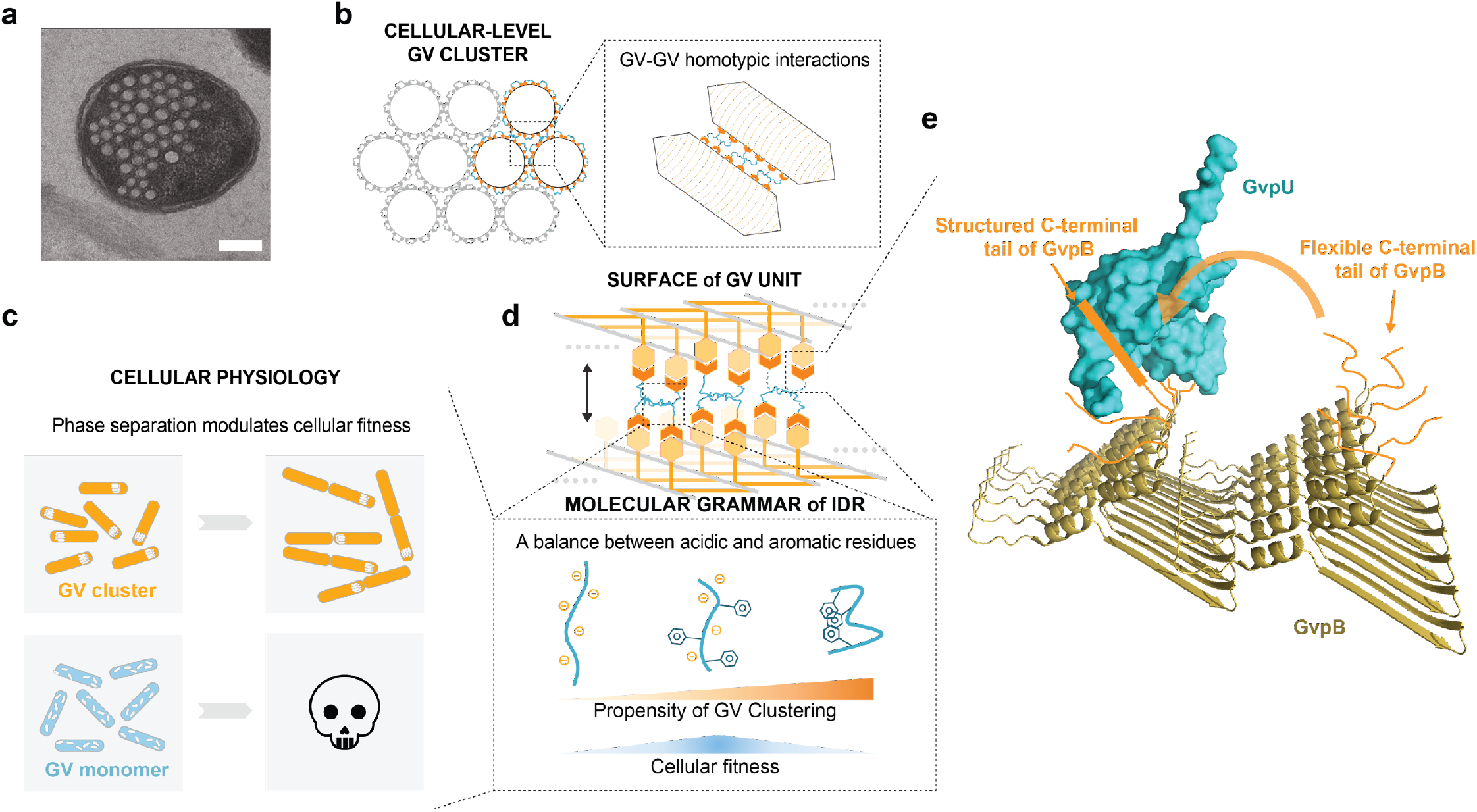
Proposed model of the assembly of GV cluster, its relation to cellular fitness, and a structure-based model of GvpU-GvpB interaction. **a**, Thin-sectioned TEM image of an *E. coli* cell containing hexagonally packed, clustered GVs. Scale bar = 200 nm. **b**, GV-GV homotypic interactions trigger the GV cluster at the cellular level. **c**, The balance between acidic and aromatic residues on IDR mediates a propensity of GV clustering and modulates cellular fitness. **d**, A model summarizing how phase separation modulates cellular fitness. **e**, Schematics of a structure-based mechanism of GvpU interaction with GvpB. The structured region of GvpB determined by cryo-EM is colored in gold, an AlphaFold model structure of GvpU in cyan, and the C-terminal tail of GvpB in orange. The C-terminal tail is drawn as a flexible coil when GvpU is not bound, and as a bar that represents a structured domain when GvpU is bound.

Lastly, although GVs have been repurposed for a range of biomedical applications, the protein engineering of GVs was thus far limited to the modulation of their mechanical and acoustic properties, and our work has added a new dimension of controlling the high-ordering clustering of these nanostructures, which can strongly influence imaging contrast, biodistribution, and cellular interactions. Future studies could leverage the genetically encodable clustering control method developed here for various applications. For example, toggling between clustered and unclustered form in response to a metabolite or cellular signal is a common method to design imaging-based biosensors, as demonstrated in the pioneering work of using the aggregation of superparamagnetic nanoparticles to sense protein-protein and protein-DNA interactions^44^. In most of the GV-based imaging methods including ultrasound, MRI and optical coherence tomography, toggling between clustered and unclustered states can produce a large difference in image contrast. Thus, linking the clustering control with protein or metabolites of interest will lead to a biosensor design that can be accessed non-invasively in deep tissues. In sum, we anticipate our studies will open doors to a wide range of future studies on both the fundamental understanding and biological applications of this unique class of protein organelles.

## Acknowledgements

We thank the Shared Equipment Authority (SEA) at Rice University for the access to core facilities and instruments, and Dr. Wenhua Guo for the training. We thank the Thyer Lab at Rice University for providing template plasmids encoding the spectinomycin resistance gene and the p15A and CloDF13 origins of replication. This work was supported by the Cancer Prevention and Research Institute of Texas (CPRIT), the NIH (R00 EB024600 and R21 EB033607), the Welch Foundation, G. Harold and Leila Y. Mathers Foundation, Hearing Health Foundation, and John S. Dunn Foundation. M.I. acknowledges support from German Research Foundation (DFG) Postdoctoral Fellowship, and Z.L. acknowledges support from Edgar O’Rear and Mary F.D. Morse Travel Award from the Institute of Biosciences and Bioengineering at Rice University. This work was supported by the Air Force Office of Scientific Research (FA9550-20-1-0241 to L.Y, A.C).

## Author contributions

Conceptualization, G.J.L., A.C., L.Y., Z.L., Q.S., and Y.D.; Methodology, G.J.L., A.C., L.Y., Z.L., Q.S., and Y.D.; Investigation, Z.L., Q.S., Y.D., A.P.A., M.I., R.L., B.Z., and M.D.M.; Formal Analysis, Z.L., Q.S., Y.D., A.P.A., M.I., R.L., B.Z., and M.D.M.; Writing – Original Draft, G.J.L., A.C., L.Y., Z.L., Q.S., and Y.D.; Supervision and Funding Acquisition, G.J.L., A.C., L.Y..

## METHOD DETAILS

### Cloning, expression, and purification of GVs

The plasmid pST39-pNL29 that encodes GV gene clusters from *B. megaterium* and the plasmid ARG1 that encodes GvpA from *A. flos-aquae* were obtained from Addgene (#91696 and #106473). Oligos for molecular cloning were synthesized (Integrated DNA Technologies (IDT), Coralville, IA). All the GV variants including those based on GvpA were cloned into pST39 vector backbone using Gibson Assembly (New England Biolabs (NEB), Ipswich, MA). For GV expression, plasmids were transformed in BL21 Star™ (DE3)pLysS One Shot™ *E. coli* strain (Thermo Fisher Scientific, Waltham, MA), which were then cultured in LB Miller Broth (Thermo Fisher Scientific, Waltham, MA) containing 100 μg/ml Carbenicillin (Gold Biotechnology, Olivette, MO), 25 μg/ml Chloramphenicol (MilliporeSigma, Burlington, MA) and 0.2% glucose (MilliporeSigma, Burlington, MA). Cells were grown at 30 °C until the optical density at 600 nm (OD_600_) reached 0.4, and then induced with 20 μM Isopropyl β-d-1-thiogalactopyranoside (IPTG) (Teknova, Hollister, CA) for 22 hrs at 28.5 °C. Cells were collected and pelleted by centrifugation at 400 x g in 50 mL conical tubes for 8 hrs with a liquid depth ≤ 10 cm. The middle layer between the buoyant cells and the sedimented cells was removed and the remaining cells were mixed with 4 mL SoluLyse-Tris (Genlantis, San Diego, CA) per 50 ml culture and 250 μL/mL lysozyme (MilliporeSigma, Burlington, MA) and 10 μL/mL RNase-free Deoxyribonuclease I (DNase I, MilliporeSigma, Burlington, MA). After 30 min rotation at 4°C, the lysate was transferred to 2mL tubes and centrifuged for 2 hrs at 400g at 4°C. The buoyant gas vesicles (GVs) located at the top of the liquid surface were isolated by removing the liquid and cell pellets underneath them using a syringe. The isolated GVs were then washed by resuspending them in 1X PBS (Teknova, Hollister, CA) followed by centrifugally-assisted flotation for three cycles.

### Cloning, expression, purification, and characterization of GvpU protein

The open reading frame of *gvpU* and *gvpU::sfGFP* were cloned into the pST39 vector with a C-terminal 6 x His-tag. The transformed cells were grown at 30°C until OD_600_ reached 0.6 and then induced with 400 μM IPTG for 20 hrs at 18°C. Cells were harvested, resuspended in 20 mM Tris-HCl lysis buffer, and lysed by 5 cycles of freeze-thaw followed by 5 cycles of probe sonication. DNase I, Benzonase^®^ Nuclease (MilliporeSigma, Burlington, MA) and Dithiothreitol (DTT) were added at 20 μg/mL, 5000 units/mL, and 1 mM, respectively, followed by 30 min incubation. Next, the samples were centrifuged for 30 min at 24,000 x g and the supernatant was loaded into a Ni-NTA affinity chromatography (Qiagen USA, Germantown, MD). After washing at 20 mM imidazole, the target protein was eluted with 300 mM imidazole. Fractions containing the target protein were collected and stored at –80°C. The purity of GvpU was verified by SDS-PAGE from the FluorChem M imager, and matrix-assisted laser desorption/ionization time-of-flight mass spectrometry (Bruker AutoFlex Speed MALDI ToF, Bruker) was used to confirm the identity of GvpU.

### Transmission electron microscopy

For negatively stained TEM of purified GVs, samples were diluted to OD_500_ = 0.2 in 1 X PBS and loaded to 200-mesh carbon-coated copper grids (Ted Pella, Redding, CA) for 3 minutes. Excess liquid was carefully blotted away with filter paper. The samples were then stained with a 2% (w/v) solution of uranyl acetate (Electron Microscopy Sciences, Hatfield, PA). High-resolution TEM images were captured using a JEOL JEM-2010 TEM and JEOL JEM-2100 Field Emission Gun TEM. For thin-sectioned TEM of GV-expressing cells, cells were fixed overnight in Karnovsky’s fixative (Electron Microscopy Sciences, Hatfield, PA), post-fixed for one hour in 1% osmium tetroxide, dehydrated in a graded series of ethanol, embedded in epoxy resin, and polymerized overnight at 70 °C. Ultra-thin sections of 100 nm thickness were cut using an ultramicrotome (Leica EM UC7), placed on an unsupported 200 mesh copper grid, and stained with uranyl acetate and lead citrate. Images were collected using a JEOL JEM-1400Flash at 120 kV equipped with an AMT NanoSprint15 sCMOS sensor.

### Hydrodynamic size measurement

The hydrodynamic size (diameter) of clustered and un-clustered GVs was measured on Malvern Zen 3600 Zetasizer (Malvern, UK). Purified GV samples were diluted to OD_500_ = 0.2 in 1X PBS and 500 μL of each sample was transferred into the cuvette (Thermo Fisher Scientific, Waltham, MA). A minimum of three measurements per sample were taken and each type of GV had at least 3 biological replicates.

### *In vitro* reconstitution of GV clusters by purified GvpU

For the reconstitution of purified GvpU to urea-treated wildtype (ureaWT), *BΔgvpU*, or *AΔgvpU* GVs described in **Fig. 2b, 2c**, and **2d**, 500 μL of GVs with OD_500_ = 0.2 were mixed with 100 μL of 30 μM purified GvpU. The molar ratio is 1.98 and 6.17 for GvpU:GvpB and GvpU:GvpA, respectively, calculated by the previously described molecular weight of GVs ^24^. For experiments described in **Fig. 2e** and **2f**, GvpB or GvpA-based GVs were purified and OD_500_ was adjusted to 0.5. Purified GvpU at 0.5 mg/mL was then added to the GV samples at a 1:5 v/v ratio (GvpU:GV). After 30 minutes of incubation at room temperature, the GVs were purified by centrifugally-assisted flotation, washed once with 1X PBS, and then dissolved in neat formic acid at a 1:9 volume ratio. The bottom fraction of the mixture was also collected. All samples were lyophilized overnight prior to sample preparation for SDS-PAGE analysis. The reconstitution experiments of ureaWT GVs described in **Fig. 6g** and **6h** were carried out by resuspending GVs in 25 mM Tris-HCl, 150 mM NaCl and 10% w/v 1,6-Hexanediol (1,6-HD, Thermo Fisher Scientific, Waltham, MA) or 12 mM adenosine triphosphate (ATP, Thermo Fisher Scientific, Waltham, MA). The final OD_500_ was set to 0.2. Purified GvpU in 25 mM Tris-HCl and 150 mM NaCl were titrated into GV samples with final concentrations listed in the figure. The hydrodynamic size (diameter) of the GVs after each titration was determined using the method described above.

### Structure prediction and docking analysis

The structures of GvpU and GvpB in **Fig. 3** were predicted using the ROSETTA online protocol ^45^. The GvpB assembly was generated as GvpB-GvpB-docking using the ClusPro web server (https://cluspro.org) following protocols ^46-49^. The choice of the simulated model was based on previously published literature on GV shell proteins ^50, 51^, and GvpU and the assembled GvpB were docked using ClusPro with the same protocol. The top 10 models with the highest balanced coefficient scores were selected and the binding sites were analyzed using PyMOL (The PyMOL Molecular Graphics System, Version 2.0, Schrödinger, LLC). The structure of GvpU in **Fig. 7** was computed on the Google Colab cloud interface using the AlphaFold framework ^52, 53^. Multiple sequence alignments (MSAs) were generated using MMseqs2 and the default setting, and the analysis resulted in five models that were sorted by their pLDDT score, from which the top-ranked model is selected. The structure of GvpB was from the cryo-EM structure (Protein Data Bank: 7R1C) ^27^.

### Pull-down assay

GvpU was fused with a C-terminal GS linker and 6x HisTag and used as the bait protein, and fragments of GvpA or GvpB were linked to the C-terminus of *Superfolder* GFP (sfGFP) as the prey proteins. The cloning, expression, and cell lysis follow the same procedures as the purification of GvpU, and the pull-down assay follows the protocol provided by Pierce™ His Protein Interaction Pull-Down Kit (Thermo Fisher Scientific, Waltham, MA). The fluorescence intensity (excitation = 485, emission = 510) of 150 μL prey proteins in 96 well plates (Corning® 96 Well CellBIND® Microplates, Corning) using a microplate reader (Agilent BioTek Synergy H4 Hybrid, Agilent). Pulled-down percentages were normalized as

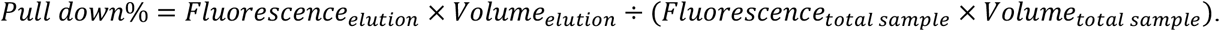

### Hydrostatic collapse pressure measurements

The apparatus for pressurized absorbance spectroscopy follows the previously described setup ^24^. Purified GVs were diluted to OD_500_ between 0.2 and 0.3 in 1 X PBS and loaded into a pressurizable cuvette (176.700-QS, Hellma GmbH & Co. KG, Müllheim, Germany). OD_500_ was measured by a spectrophotometer (STS-VIS, Ocean Optics, Dunedin FL, USA), while the cuvette was pressurized by a single-valve pressure controller (PC series, Alicat Scientific, Tuscon, AZ, USA) from 0 to 1.4 MPa at 0.2 MPa per step.

### Phase separation assay of purified GvpU and GvpU::sfGFP

Purified GvpU was dialyzed using 25mM Tris-HCl buffer containing 500mM NaCl and then concentrated using an Amicon® Ultra-15 Centrifugal Filter Concentrator (MilliporeSigma, Burlington, MA) with a 10kDa molecular weight cut-off. The resulting sample was centrifuged for 5 minutes at 13,000 x g to collect the supernatant, which contained GvpU in the dilute phase. The concentration of GvpU was then measured using absorbance at 280 nm (OD_280_) on a Take3 plate (Agilent BioTek Synergy H4 Hybrid, Agilent) and adjusted to a final concentration of 1 mM. 20 μL of the GvpU solution was loaded onto a cover glass petri dish (MatTek Corporation, Ashland, MA), and a Nikon Eclipse Ti2 (Nikon) fluorescence microscope was used for imaging with a 10x objective in bright-field mode with phase ring 1. To induce phase separation, 20 μL of 25 mM Tris-HCl buffer containing 0 mM NaCl was added to the GvpU droplet on the cover glass. The camera was set to take an image every 10 seconds, starting immediately after the addition of the NaCl-free buffer.

The same dialysis, concentration, and concentration measurement procedures were used for purified GvpU::sfGFP, with the only difference being the final concentration of NaCl in the buffer. 20 μL of 0.9 mM GvpU::sfGFP was loaded onto the cover glass and serially diluted to the concentrations listed in **Fig. 5c**. For imaging, the Nikon Eclipse Ti2 (Nikon) fluorescence microscope was switched to fluorescence mode with a 10 x objective and the emission wavelength was set to 515 nm to detect the sfGFP. Images were taken for each protein concentration with the exposure time kept at 100 μs.

### Sedimentation assay

A sedimentation assay was used to measure the binodal of GvpU as described previously^54^. Briefly, protein samples were prepared at a total concentration of 150 μM in 25 mM Tris-HCl buffer with 150 mM NaCl and incubated at different temperatures for 2 h. The dilute and dense phases were then separated by centrifugation at the same temperature as indicated. The dilute-phase concentration (the left binodal) was measured on a Nanodrop Microvolume spectrophotometer (Fisher Scientific) by assessing OD_280_, and the concentration was calculated from the extinction coefficient of GvpU. To measure the dense-phase concentration, 5 μL of the separated dense phase was removed using a positive displacement pipette (RAININ) and diluted into 95 μL of buffer containing the same concentration of salt as the sample with the addition of 6 M urea. Different dilution ratios were applied to ensure measurement accuracy. The dense-phase concentration (the right binodal) was measured by OD_280_ on a Nanodrop and calculated based on the dilution ratio and the extinction coefficient.

### Temperature-dependent turbidity measurement

The temperature-dependent turbidity test of GvpU was performed on a Cary 300 temperature-dependent ultraviolet-visible spectroscopy equipped with a multicell thermoelectric temperature controller. The samples of GvpU were prepared in 25 mM Tris-HCl and 150 mM NaCl. A cooling rate of 1 °C/min was applied to the samples while the absorbance at 350 nm was recorded at every 0.33 °C increment. Normalized turbidity was calculated by the absorbance at the highest temperature point divided by the absorbance at the lowest temperature point.

### Live-cell fluorescence microscopy

*E. coli* cells were grown as described above. At different time points after IPTG induction, cells were pelleted and washed 3 times with Hanks balanced salt solution (HBSS) (ThermoFisher). Cells were diluted back into HBSS and placed onto a 35-mm number 1.5 coverslip glass-bottom dish. A Leica Stimulated Emission Depletion (STED) SP8 (Leica) confocal microscopy was used for imaging with a ×100/1.40-oil immersion objective. The white light laser (WLL) power source was set at 70% with emission intensity at 2% and emission wavelength at 585 nm for mCherry. The Leica Hybrid detector was set at 590 nm to 660 nm.

### Cellular fitness tracking

*E. coli* cells containing the various GV variants were grown at 30 °C until they reached OD_600_ of 0.4. The cells were then induced with 20 μM IPTG for 5 hrs at 28.5 °C. Afterwards, the cells were transferred to a new 50 mL LB culture without IPTG at an initial OD_600_ of 0.05. The growth of the new culture was monitored every hour by measuring OD_600_.

### Quantification and statistical analysis

The statistical information of the experiments, including sample size (n), and P-values, can be located in the figures, captions for the figures, and the method section. The statistical analysis was carried out using GraphPad Prism software and is represented as the mean ± standard deviation (STD). For all the multiple comparisons, the Welch and Brown-Forsythe ANOVA tests were performed, which is a version of one-way ANOVA that does not assume that all the groups were from a population with equal variances. p<0.05 was set as significantly different.

