## Supplementary Material for "Spatial Organization of Gas Vesicles is Governed by Phase-separable GvpU"

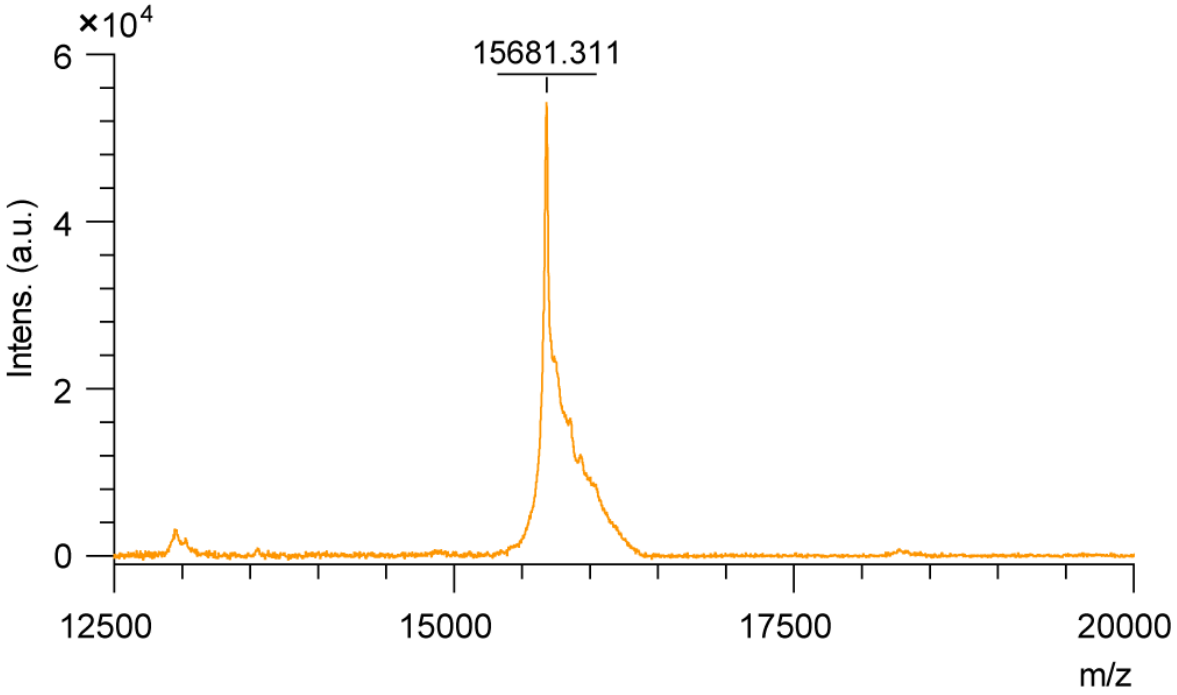


**Figure S1. Mass spectrum of GvpU::6xHisTag obtained from MALDI TOF MS.** The x-axis displays the mass-to-charge ratio (m/z) of the ionized species, while the Y-axis represents the ion intensity. The presence of the peak with 15681.311 m/z in the spectrum confirms the identity of GvpU::6xHisTag.

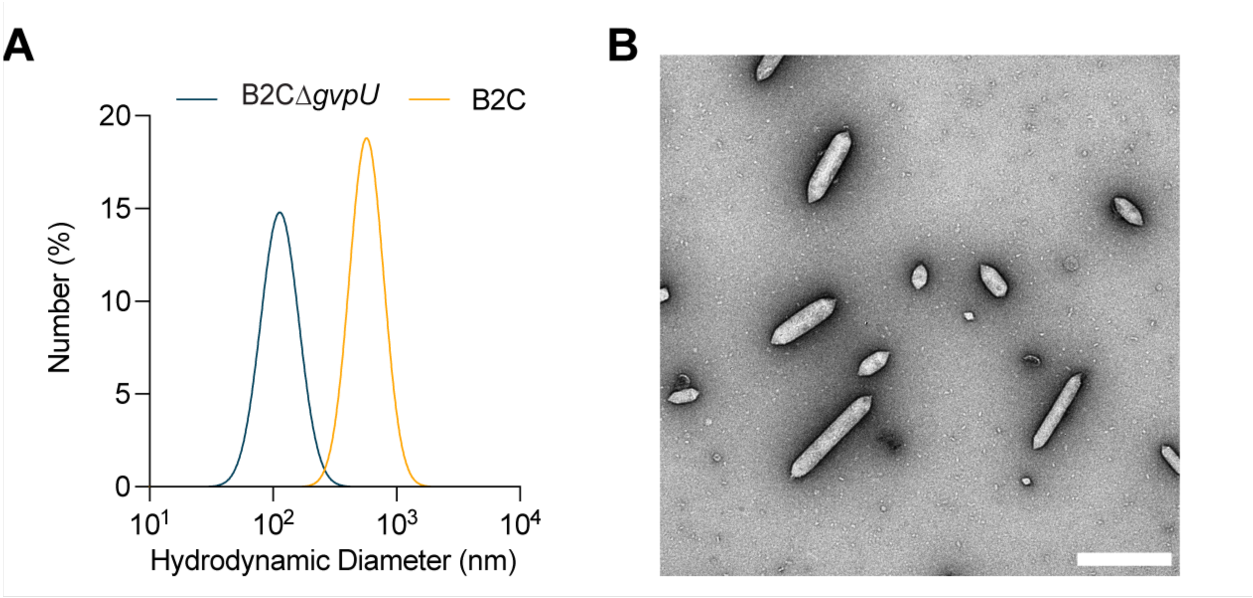


**Figure S2. GvpC does not interfere with GvpU-mediated GV clustering.**

(A) Representative DLS measurement of *B2CΔgvpU* constructs (n = 3, biological replicates).

(B) Representative TEM images of *B2CΔgvpU* constructs (scale bar, 500 nm).

**Table S1 Detailed amino acid sequence of GvpA and GvpB fragment.**

| Name | Sequence Range | Sequence |
|---|---|---|
| $A_{(Nt)}$ | M1-S10 | MAVEKTNSSSS |
| $B_{(Nt)}$ | M1-S10 | MSIQKSTNSS |
| $A_{(\alpha1)}$ | A13-G23 | AEVIDRILDKG |
| $B_{(\alpha1)}$ | A13-G23 | AEVIDRILDKG |
| $A_{(\beta1)}$ | V25-L34 | VIDAWVRVSL |
| $B_{(\beta1)}$ | V25-V34 | VIDAFARVSV |
| $A_{(\beta2)}$ | E38-S50 | ELLAIEARIVIAS |
| $B_{(\beta2)}$ | E38-S50 | EILTIEARVVIAS |
| $A_{(\alpha2)}$ | V51-V60 | VETYLKYAEAV |
| $B_{(\alpha2)}$ | V51-V60 | VDTWLRYAEAV |
| $A_{(Ct)}$ | T64-A71 | TQSAAVPA |
| $B_{(Ct)}$ | L64-I88 | LRDDVEENGLPERSNSSEGQPRFSI |
